# Cerebrospinal fluid and blood profiles of transfer RNA fragments show age, sex and Parkinson’s disease-related changes

**DOI:** 10.1101/2022.06.29.498078

**Authors:** Iddo Paldor, Nimrod Madrer, Shani Vaknine-Treidel, Dana Shulman, David S Greenberg, Hermona Soreq

**Affiliations:** The Neurosurgery department, Rambam Health Care Campus, Haifa; The Edmond and Lily Safra Center for Brain Sciences, The Hebrew University of Jerusalem; Department of Biological Chemistry, The Alexander Silberman Institute of Life Sciences, The Hebrew University of Jerusalem; The Rachel and Selim Benin School of Computer Science and Engineering, The Hebrew University of Jerusalem, 9190401 Jerusalem, Israel

## Abstract

Transfer RNA fragments (tRFs) have recently been shown to be an important family of small regulatory RNAs with diverse functions. Recent reports have revealed modified tRF blood levels in a number of nervous system conditions including epilepsy, ischemic stroke and neurodegenerative diseases, but little is known about tRF levels in the cerebrospinal fluid (CSF). To address this issue, we studied age, sex and Parkinson’s disease (PD) distributions of tRFs in the CSF and blood data of PD patients and healthy controls from the NIH and the PPMI small RNA-seq datasets. The higher levels of long tRFs were found in the CSF than in the blood. Furthermore, the CSF showed pronounced age-associated declines of the level of 3’-tRFs and i-tRFs and more pronounced differences between the sexes. Blood showed moderate elevation of 3’-tFs levels with age. In addition, different distinct sets of tRFs segregated PD patients from controls in the CSF and in the blood. Finally, we found enrichment of tRFs predicted to target cholinergic mRNAs (Cholino-tRFs) in the mitochondrial originated tRFs, raising the possibility that the neurodegeneration-related mitochondrial impairment may lead to deregulation of cholinergic tone. Our findings suggest that CSF expressed tRFs are not a mirror of blood tRFs but rather potentially reflect the cerebral changes. Further, both CSF and blood present modified levels of tRFs in a sex-, age-and disease-related manner, calling for including this important subset of small RNA regulators to future studies.

## Introduction

### The cerebrospinal fluid (CSF) flows in a closed circuit

CSF is produced from blood in the choroid plexus, which is located in the cerebral ventricular system, and supplied by different choroidal arteries (Brodbelt and Stoodley 2007; Sakka *et al*. 2011; Li *et al*. 2019; Orešković *et al*. 2017). CSF then flows through the ventricular system - lateral ventricles, third ventricle and fourth ventricle. After exiting from the lateral and medial foramina of the fourth ventricle, the fluid surrounds the brain and the spinal cord in the subarachnoid space throughout the central nervous system (CNS) (Johnston and Teo 2000; Brandner *et al*. 2014; Djukic *et al*. 2016). This circle ends with reabsorption of the CSF to the blood stream by the specialized arachnoid granulations, adjacent to the large venous sinuses around the brain (Wichmann *et al*. 2021; Proulx 2021; Chen *et al*. 2015).

Apart from providing a physical “shock absorber” for the brain, the CSF also nourishes the brain with nutrients, and serves to remove metabolic byproducts from the brain (Spector *et al*. 2015; Herrera *et al*. 2018; Tumani *et al*. 2017). During the production of CSF, certain elements – cells, electrolytes and proteins – are actively excreted into the CSF, while others are filtered so that they are decreased, or nonexistent, in the CSF. Thus, when compared to peripheral blood, the levels of some molecules are unchanged while others are elevated or decreased. Although difficult to document, it is important to note that upon its return to the bloodstream, the CSF contains both soluble components that originate in the blood and soluble ones that originate in the neurons and glia of the CNS.

Protein levels are significantly lower in the CSF than in the blood. The normal level of protein in the CSF is about 35 Mg/dL, compared to 7000 in the peripheral blood. Elevated levels of protein are hence indicative of a pathological process (McCudden *et al*. 2017; Hühmer *et al*. 2006; Brooks *et al*. 2019). After production from the blood, CSF courses through the central nervous system (CNS). During this course, and prior to reabsorption, other elements are secreted into the fluid by the nervous and glial elements that surround it. Therefore, there is certainly a gradient of concentrations along the course of CSF flow. Although difficult to document, it is important to note that upon its return to the bloodstream, the CSF contains both soluble components that originate in the blood and soluble ones that originate in the neurons and glia of the CNS. In the peripheral nervous system (PNS), neuronal acetylcholine (ACh) participates in sympathetic and parasympathetic peripheral signaling (Soreq 2015). In the neuro-immune interface, ACh signaling is elevated under anxiety and can also function as an immune modulator with the ACh anti-inflammatory response suppressing inflammation (Shaked *et al*. 2009). Taken together, these facts call for in-depth testing of the cholinergic deficits in sncRNA regulators of cholinergic network genes whose functioning is altered in both the brain and the CSF.

### CSF sampling methods matter

Comparing CSF taken from the lumbar cistern through a lumbar puncture to CSF taken from the lateral ventricle revealed differences in the concentration of diverse compounds. Correspondingly, exogenous drugs given, which are all blood-derived, have a much higher concentration in ventricular CSF compared to lumbar CSF (Bohn *et al*. 2019). For instance, the concentration of zidovudine and lamivudine (both anti-HIV medications) was shown to be up to four times greater in ventricular CSF compared to lumbar sampling (Eggers *et al*. 2020). Further, elements that originate from the brain, such as Tau protein and neuron-specific enolase are present in higher concentrations in CSF taken from the ventricle. In contrast, there is a higher level in the lumbar CSF of compounds that are derived from the meninges, such as cystatin C and beta-trace protein (Wajda *et al*. 2020). It seems that the order of tissues which contribute to CSF composition is peripheral blood first, then brain and then meningeal contribution. Thus, elements in CSF that are derived from the blood decrease throughout the course of CSF flow, whereas CNS-derived elements increase throughout the course of this flow.

### Small Non-coding RNAs are important but understudied regulators of brain functioning

aThe great majority (∼ 98%) of transcripts of the human genome are non-coding RNAs (ncRNAs). ncRNAs are encoded by a highly diverse class of genes and their length ranges from dozens to tens of thousands of nucleotides (Mattick and Makunin 2006); they are classed as either long noncoding RNA (longer than 200 nucleotides) or short/small noncoding RNAs. There are numerous classes of sncRNA including miRs, (Kleaveland *et al*. 2018; Bartel 2018) siRNAs, piRNAs, snoRNAs, snRNAs, and the newly rediscovered transfer RNA fragments (tRFs) (Pederson 2010) (Fig. 1), which may originate from nuclear or mitochondrial tRNA genes.

**Figure 1.**
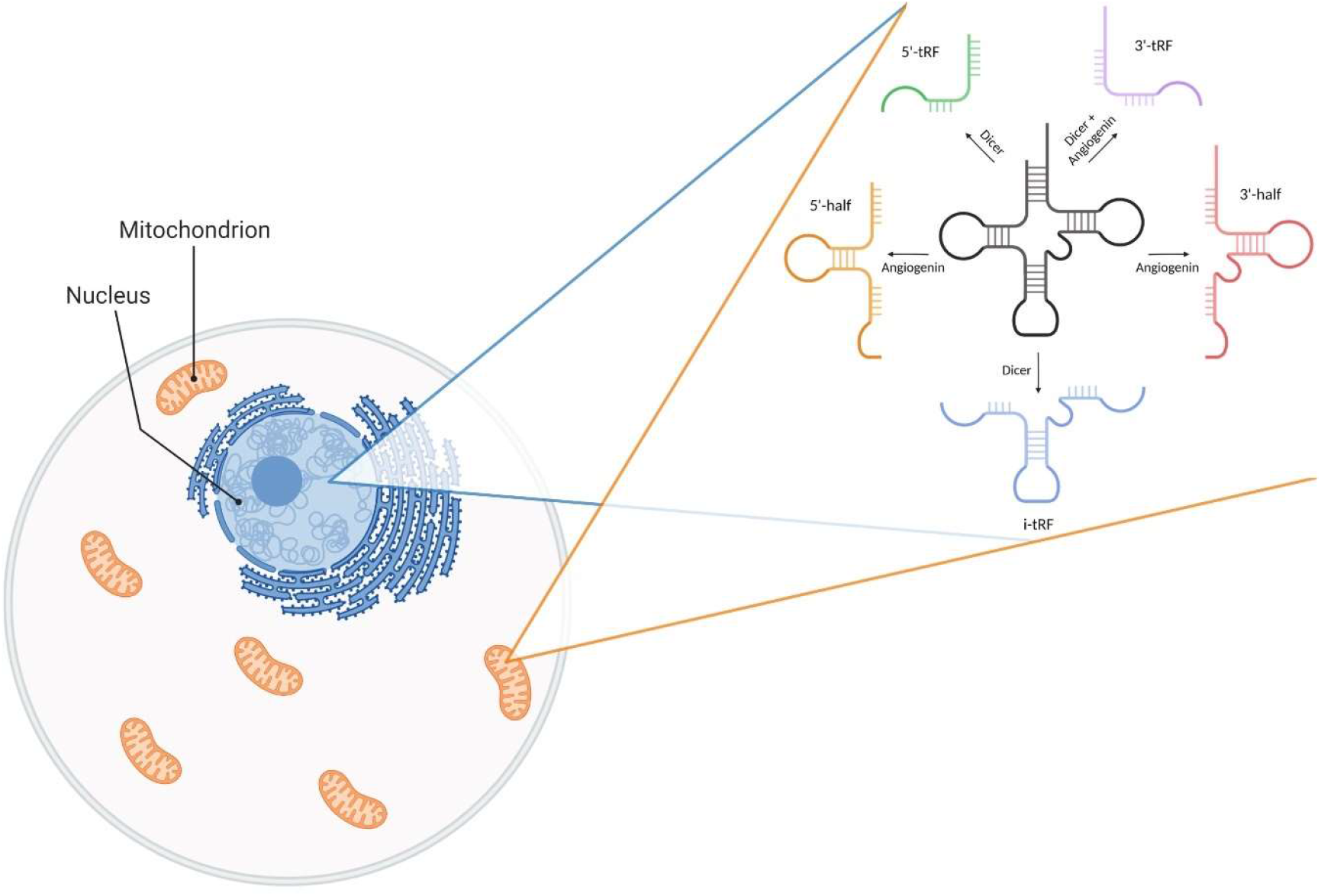
tRFs may originate from both nuclear and mitochondrial tRNA genes and are classified into several sub-groups according to the nuclease-mediated break patterns that led to their formation. Created with BioRender.com.

TRFs have been shown to play a role in cancer (Yu *et al*. 2020) and other diseases (Shen *et al*. 2018; Soares and Santos 2017), but the biological functions and role in disease pathogenesis of many of them is not well understood. In principle, sncRNAs may function within both the nervous and the immune systems as ‘negotiators’ that may communicate between these two interacting compartments by targeting regulatory genes that enable co-modulation of immune and neuronal processes (Soreq and Wolf 2011; Shaked *et al*. 2009). This would also be highly relevant for the cholinergic network, which is an important modulator of cognition and behavior, controls the complex temporal dynamics of higher brain functions (Paolone *et al*. 2013; Sarter and Bruno 1997) and may be involved in the pathogenesis of numerous nervous system diseases.

Expression of specific sncRNAs in the CSF is indicative of both disease existence and, in certain instances, of disease progression, severity and response to treatment. Specifically, tRFs rapidly emerge as an important newly re-identified family of sncRNAs capable of operating like miRs and many other functions (Winek *et al*. 2020). Therefore, we set out to seek these characteristics in healthy male and female individuals along age and with or without Parkinson’s disease (PD).

## Methods

### Data acquisition

To identify changes in the CSF’s tRF profiles, we have first analyzed a short RNA-seq dataset (phs.000727 NIH (Burgos *et al*. 2014)) of post-mortem CSF samples from 60 PD patients (23:37 females/males) and 63 age-matched healthy controls (Ctrl). Originally, the data contained 69 controls and 66 PD patients. 6 controls and 6 PD patients were removed, a priori, as outlier samples since they were apart from all patients in a PCA test.

A yet larger blood short RNA sequencing data from both PD patients and healthy controls was obtained from the Progression Markers Initiative (PPMI) database (https://www.ppmi-info.org/accessdata-specimens/download-data; updated information on the PPMI study is available at www.ppmi-info.org). Only samples of healthy controls (Ctrl) and idiopathic PD patients, who did not carry any of the known PD genetic markers, were used. For each participant the latest of the five available time points was chosen and only samples with RIN ≥ 6 were introduced into the analysis. Distribution of samples of both cohorts is available in table 1.

**Table 1.** Distribution of RNA samples across datasets, diagnosis and sex. In bracket is the average of age for each group.

| Tissue | CSF (NIH) |  | Blood (PPMI) |  |
| --- | --- | --- | --- | --- |
| Diagnosis | PD | Ctrl | PD | Ctrl |
| Male | 37 (64.99) | 36 (64.21) | 240 (78.16) | 119 (81.39) |
| Female | 23 (64.21) | 27 (62.13) | 120 (81.43) | 64 (85.41) |

### Data analyses

Short RNA samples of both cohorts were filtered and cleaned by Flexbar (Roehr *et al*. 2017; Dodt *et al*. 2012), followed by alignment to MINTbase using MINTmap (Loher *et al*. 2017; Pliatsika *et al*. 2016). Follow up analyses were done in R with count per million (CPM) normalization, with ggplot2 package (Wickham 2016) for drawing up graphs. Linear discriminant analysis (LDA) was done using MASS package (Venables and Ripley 2002) in R as well, on counts which were normalized by the independent filter of the DESeq package (Love *et al*. 2014) and filtered so only tRFs which had median ≥ 10 were kept. Influence score for each tRF was calculated by multiplying the median of their expression across all samples in their linear discriminant coefficient from the LDA analysis, separately for the two LD components.

## Results

To approach the putative role of sncRNAs in the CSF as compared to blood, we selected the family of tRFs and analyzed their subtypes, levels and content in the CSF of men and women as compared to their blood profiles. Notably, while CSF presented tRFs of most lengths and of all subtypes, the blood tRF expression was far more limited and consisted mainly of i-tRFs and 3’-tRFs. Importantly, 3’-halfs were almost completely absent in the blood. (Fig 2). Further, in the blood almost all of the expressed tRFs were of 33 bases or less across all subtypes, while in the CSF a more varied distribution was visible. (Fig.2)

**Figure 2.**
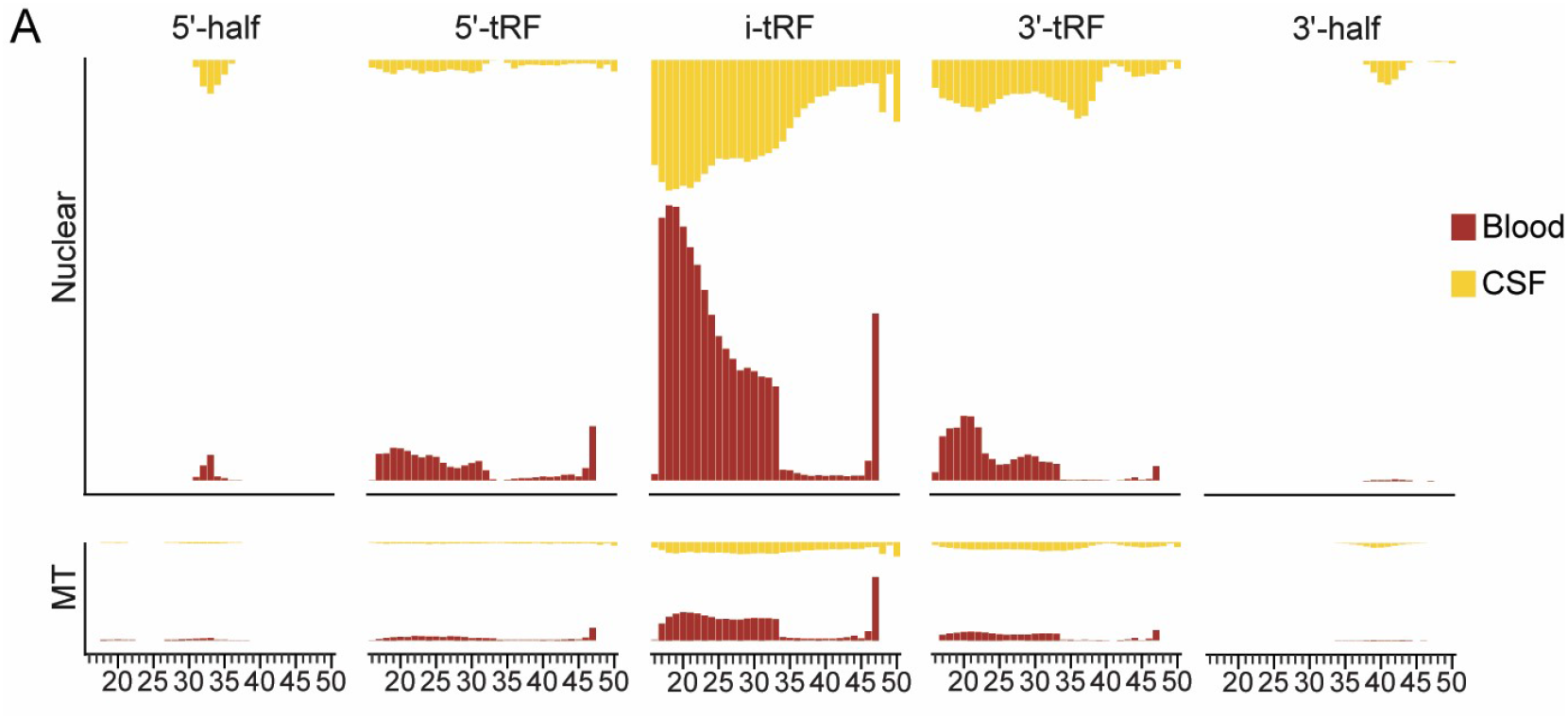
Short tRFs and half-tRFs are almost absent in blood while expressed in the CSF, with mitochondrial genome-originated tRF levels being far lower in both. Shown are the type and size distribution-based profiles of nuclear genome-originated tRFs (top) and mitochondrial genome-originated ones (bottom) in blood (ochre) and CSF samples (orange) from healthy volunteers in the PPMI dataset. Note the higher levels of i-tRFs and 3’-tRFs compared to other subtypes, the larger amounts of shorter tRFs in both of these categories and the very low levels of mitochondrial genome-originated tRFs in both CSF and blood.

### CSF tRF profiles differ from blood tRF profile

Diverse nucleases lead the production of particular tRF subtypes (Fig 1). In the CSF, the great majority of tRFs constitutes of nuclear genome-originated i-tRFs and 3’-tRFs, with the levels of mitochondrial genome-originated tRFs being far lower (Fig 2). Additionally, the shorter tRFs in these categories appear at higher levels (Fig 2). Intriguingly, the blood levels of these tRF subtypes are both lower than those of the CSF ones, but the age-related decline in their length is considerably shallower, possibly reflecting lower levels of the responsible nucleases in the blood compared to the CSF.

### 3’-tRF and i-tRF levels vary between CSF and blood

NcRNAs can operate in a sex-specific manner and may cause male-female differences in the pathogenesis of diverse disorders (Simchovitz-Gesher and Soreq 2020), but whether this relates to the CSF and blood tRF profiles as well is still unknown. To address this matter, we sought tRF distributions of CSF and blood of healthy males and females, and found major differences in the nuclear-originated tRFs seen in these sources accompanied by more limited differences in mitochondrial-originated tRFs (Fig 3A). Intriguingly, we have further observed a steep age-related decline in the 3’-tRFs and i-tRFs constituting the majority of these CSF tRFs, contrasting a very shallow age-related elevation in the blood levels of the 3’-tRFs and a yet shallower decline in the blood i-tRF levels (Fig 3B). Furthermore, we found diverse levels of the different tRF subtypes as well as male-female differences in the levels of CSF but not blood levels of 5’-half and 3’-half tRFs (Fig 3A).

**Figure 3.**
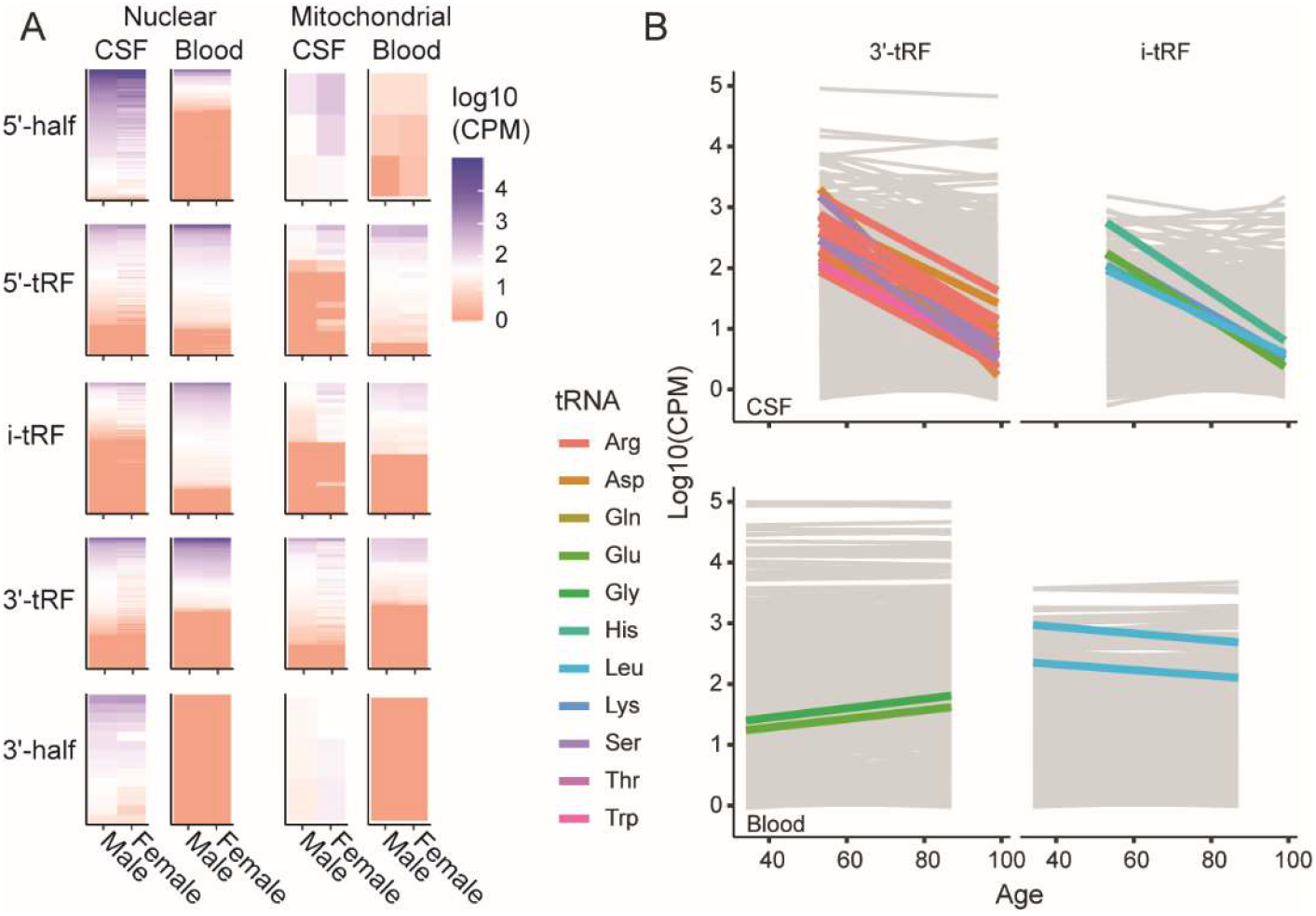
CSF and blood tRF subtypes and levels vary with sex and age. A. Shown are the levels in log10(CPM) of the diverse tRF subtypes in the CSF and blood of healthy males and females. The levels in CPM of nuclear and mitochondrial genome-originated tRFs in the CSF and blood of males and females vary between the different tRF subtypes. Also, both 5’- and 3’-halves of tRFs, and to a certain extent also 5’-tRFs show apparent differences between males and females. B. The levels in log10(CPM) of different 3’-tRFs and i-tRFs decline steeply with age in the CSF, but not blood.

### The CSF and blood tRF profiles of Parkinson’s disease patients show disease-specific patterns

To pursue the impact of disease on the tRF profiles of CSF and blood, we chose to explore the differences between PD patients and healthy controls in both the NIH and the PPMI datasets. To this end, we divided the cohorts into four groups by diagnosis and sex (PD Male, Ctrl Male, etc.) and preformed linear discriminant analysis (LDA) to study the power of tRF expression in clustering them. This analysis revealed clear clustering of PD vs. control groups, taking into account sex, which was far more clearly apparent in blood samples compared to the CSF ones (Fig. 4A, B). The separation of the clusters in the CSF plot (Fig 4A) revealed that the difference between PD and Ctrl groups manifested by LD1, whereas the separation into sex is comprised of an interaction between LD1 and LD2. In the blood (Fig 4B), LD1 separated the groups into males and females, whereas LD2 separated them into PD and Ctrl. Therefore, in the following analyses we chose to focus on LD1 in the CSF LDA and on LD2 in the Blood LDA, as both differentiate the samples according to diagnosis.

**Figure 3a.**
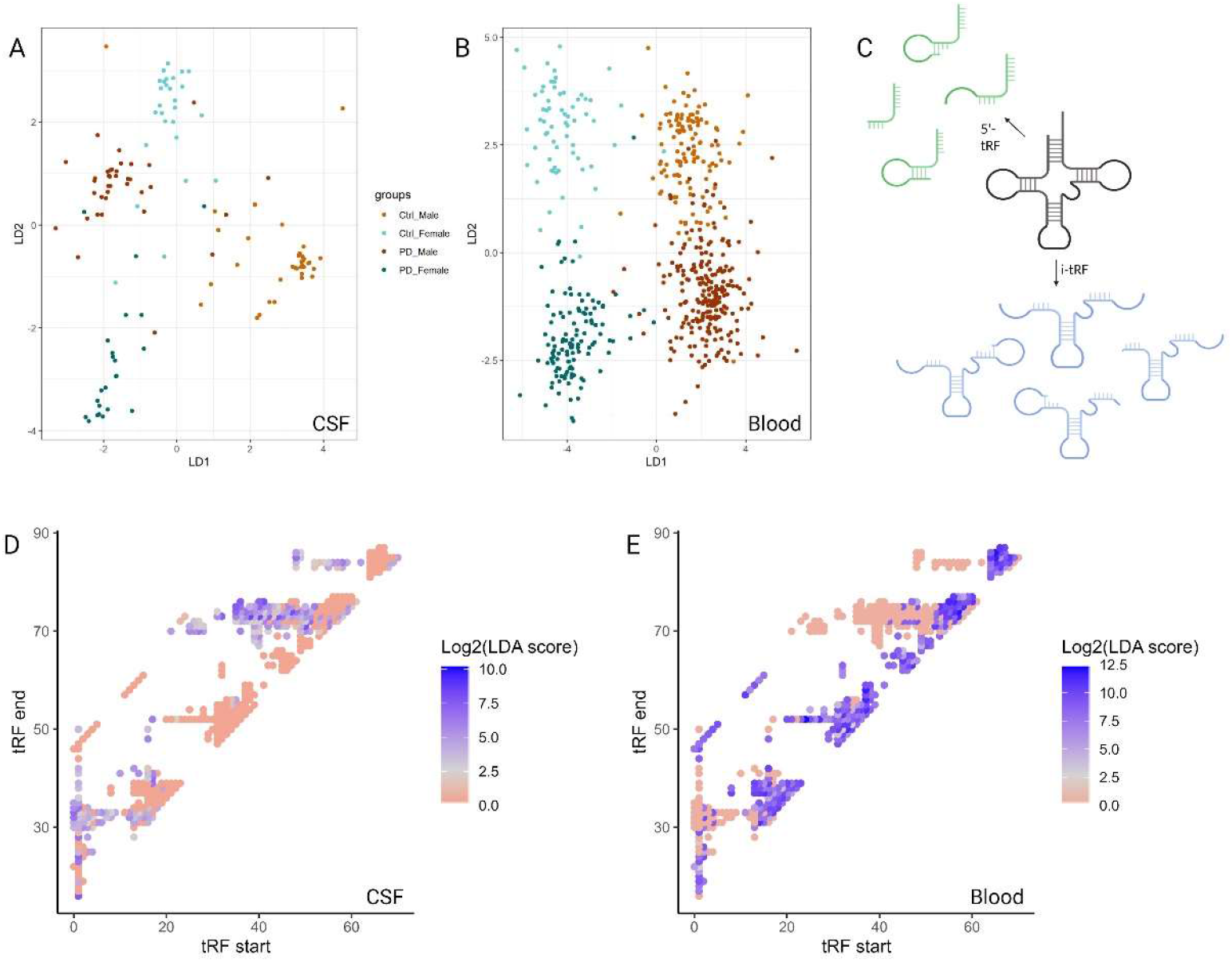
Both CSF and blood tRFs cluster male and female PD patients distinctly from controls, based on mirror pattern tRF types. Shown are the LDA segregation patterns from CSF (A) and Blood (B) of male and female PD patients and controls (see Figure for the color code). Note the more sparce segregation pattern of CSF patients and the closer packed groups of blood patients. C. Each nuclease-mediated break pattern population is comprised of tRFs of various lengths which segregate PD from controls in blood and CSF. D, E. The start and endpoints along the originating tRNA molecule of each tRF in the CSF (D) and in the blood (E). Color represents the effect of that tRF on LDA segregation of PD and control patients as indicated by the calculated influence score. Each tRF that was expressed in only one dataset received an influence score of zero in the other dataset. Created with BioRender.com.

To further zoom into the tRF profiling of CSF tRFs from PD patients and controls, we analyzed which tRFs contribute the most to the two LD components of interest, LD1 in the CSF LDA and LD2 in the Blood LDA. For each tRF we calculated the “influence score”, which comprised of a multiplication of its LD component linear discriminant coefficient by its median expression levels throughout all samples of the dataset. We then calculated log2 of the absolute value of each tRF, and plotted the resulting influence score against the start and end points of every tRF (Fig. 4D, E). Intriguingly, we found that the tRFs that affect mostly the segregation between PD and controls stem from different populations in the blood and in the CSF, reflecting distinct cut points along the originating tRNA molecules. Thus, tRFs that emerged as highly impacting the blood tRF profile had no or little impact in the CSF and vice versa, resulting in a mirror image.

### Mitochondrially originated tRFs are enriched in potential Cholino-tRFs

To examine the potential activity of tRFs as regulators of gene expression, we chose to focus on the cholinergic system that is impaired with age (Perry 1980), and in neurodegenerative diseases such as PD (Bohnen and Albin 2011). For this purpose, we sought potential targets of the observed tRFs using the MR-microT DIANA sequence-based prediction algorithm (Reczko *et al*. 2012). The likelihood of interacting with complementary binding elements in the 3’-UTR of protein coding mRNA sequences was calculated separately, creating a prediction score that was normalized by the conservation of the binding site in a number of species. Targets with prediction score < 0.8 were dropped (Fig. 5A). The specific potential of Cholinergic binding was assessed based on predicted targeting of 94 genes that have been associated with cholinergic activity in the brain, (Lobentanzer *et al*. 2019a). The genes were weighted by their direct influence on the cholinergic system (Supplementary Table X) and the cholinergic binding potential of each tRF was set as the sum of the weight of all of its predicted cholinergic targets. The threshold of enriched Cholinergic potential was set as the 85^th^ quantile of the cholinergic binding potential of all the tRFs in the CSF and blood independently (5 for both). Interestingly, while nuclear tRFs showed no specific enrichment of Cholino-tRFs, the mitochondrial originated tRFs were highly enriched with Cholino-tRFs both in the blood and the CSF (p<3×10^−17^, p<4.2×10^−7^, Fisher test, FDR; Fig. 5B). This suggests that impairment in mitochondria (which accompanies age and neurodegenerative diseases) might impact the cholinergic balance of the patient via deregulated Cholino-tRFs.

**Figure 5.**
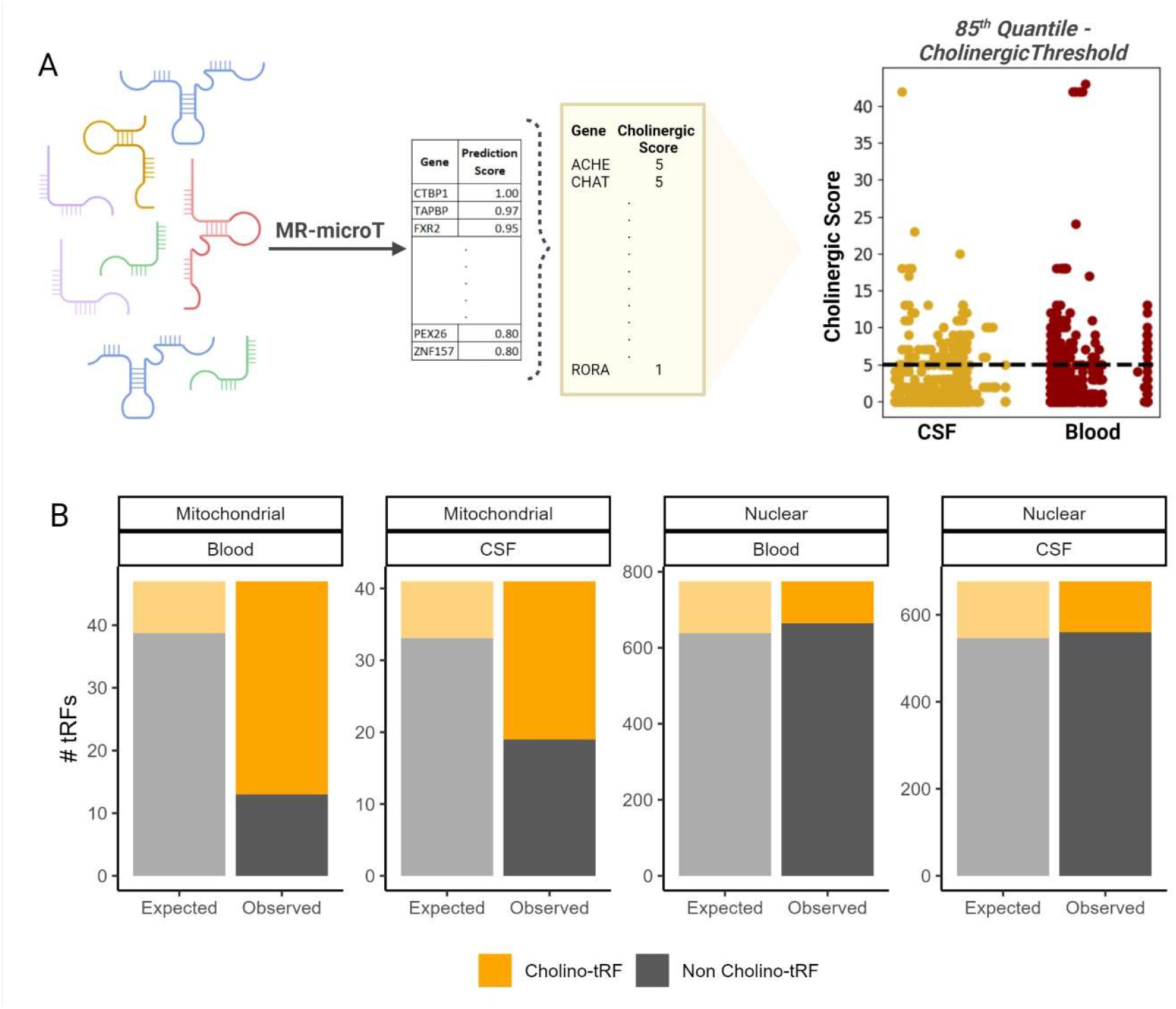
Mitochondrial tRFs are enriched with Cholino-tRFs. A. The Cholinergic potential of each tRF found in blood or CSF was set according to the number of predicted targets it had of a list of 94 known Cholinergic genes weighted according to their direct effect on the Cholinergic system. The Cholinergic score of each tRF was set as the number of Cholinergic genes multiplied by their weights, and the Cholinergic threshold was set as the 85th quantile of the Cholinergic scores in the CSF and Blood independently. B. Shown are the numbers of observed Cholino-tRFs (orange) and Non Cholino-tRFs (grey) among mitochondrial (left) and nuclear (right) tRFs in the blood and CSF, compared to their predicted share based on non-dependent distribution (expected; semi-transparent). Created with BioRender.com.

## Discussion

### CSF flow and composition may reflect age, sex and disease processes

There is a shift in CSF flow which is demonstrable during aging. The shift has been clearly shown in both sampling and imaging research(Balédent *et al*. 2004; Schmid Daners *et al*. 2012; Stoquart-ElSankari *et al*. 2007). There is growing evidence that CSF is also altered with aging in terms of its content, and in many of the elements mentioned above. CSF protein levels are clearly elevated in older age, and it has been suggested that the normal value for CSF protein be elevated in older patients(Breiner *et al*. 2019; Hegen *et al*. 2021). A similar paradigm shift appears in normal values of CSF cell composition. For instance, the yield of cells for the diagnosis of meningitis is significantly better in older patients as compared to younger ones(Mentis *et al*. 2017). It is also clear that the composition of different RNA fragments is not constant in different aging phases – in health and in disease (Osgood *et al*. 2017; Van Harten *et al*. 2015). However, it is unclear whether the changes in tRF levels in the CSF are related to the changes in flow, or occur independently from the presumed slower flow. Similarly, differences in the composition and flow of CSF have been shown between the two sexes.

Detailing the differences between male and female CSF is beyond the scope of this article. However, to note a few differences or similarities: Females have been shown to have, in average, a lower total volume of intracranial CSF, and to have a lower level of total protein in the CSF, although resting normal glucose in the CSF seems to be equal in both sexes. Finally, RNA levels have been shown to be different in the different sexes in various pathological states (Leen *et al*. 2012; Hegen *et al*. 2014; Castellazzi *et al*. 2020; Meixensberger *et al*. 2020; Teasdale *et al*. 1988; Zheleznyakova *et al*. 2021; Dangla-Valls *et al*. 2017a; Gui *et al*. 2015; Otake *et al*. 2019). Both aging and systemic diseases notably involve shedding of protein, nucleic acids and other factors into the blood. In these pathologies, the contents of CSF may be partially derived from elements that are passed on from the blood during the production of CSF. We acknowledge that there is certainly a bias regarding CSF in systemic diseases, since CSF sampling is only performed in clinical practice when there is suggestion of CNS involvement. Hence, lumbar puncture is not performed on patients with systemic disease and no signs of nervous system involvement. Therefore, it is difficult to discern the origin of elements in the CSF in patients with non-brain-specific diseases due to this bias. However, in brain-specific disease entities with no known systemic involvement, differentially expressed elements must be thought of as originating from the brain, and shed into the CSF.

Recent reports of miRs as well as tRFs regulating both neuronal and immune cholinergic processes suggest a therapeutic potential for manipulating these ncRNAs in cholinergic-related diseases, that affect both brain function and the immune system such as AD (Lau and de Strooper 2010), Parkinson’s disease (PD) (Cirnaru *et al*. 2021), epilepsy (Reschke and Henshall 2015) and anxiety-related disorders (Penner-Goeke *et al*. 2019). Specifically, sncRNAs in the CSF have been shown to be indicative of certain cerebral pathological conditions. For instance, miR levels have been shown to be altered in multiple sclerosis (MS), Alzheimer’s disease (AD), Parkinson’s disease (PD), prion diseases and amyotrophic lateral sclerosis (ALS) (Freiesleben *et al*. 2016; Saucier *et al*. 2019; Wang *et al*. 2011; Derkow *et al*. 2018; Wallach *et al*. 2021).In the case of MS, a significant difference has been shown between blood and CSF. Specifically, Zheleznyakova et al. compared the expression levels of miRs and other sncRNAs in the blood and in the CSF of patients with different forms of MS. They showed that many MS-related sncRNAs were increased in the CSF compared to non-MS controls. Conversely, the vast majority of sncRNAs in the blood mononuclear cells were **decreased** in MS patients compared to non-MS controls (Zheleznyakova *et al*. 2021).

Regarding AD, there is ample evidence of sncRNA shedding into the CSF of patients with AD. A number of miRs, such as miR-222 and the acetylcholinesterase-targeting miR-125b (Lobentanzer *et al*. 2019b), have been shown to be differentially expressed (DE) in the CSF of AD patients (Dangla-Valls *et al*. 2017b). Finally, DE sncRNAs have also been shown in the CSF of PD patients, where hsa-mir-626, for example has been the main focus – both as a potential biomarker for disease severity and as potential therapeutic targets (Qin *et al*. 2019). Thus, DE sncRNAs in the CSF should be considered representative of the brain pathology, and not of changes that were shed into the CSF during production of CSF from the blood. In this context, the currently presented analyses and datasets may serve as a Resource to enable interested researchers to explore relevant CSF and blood tRFs in the context of their sex, age and disease-related values.

## Acknowledgments

This work was supported by a grant from the Israel Science Foundation (ISF; to H. Soreq). The authors wish to thank to the US friends of the Hebrew University of Jerusalem and to Leopold Sara and Norman Israel fund, Sphardi Jews aged home for the grants for N. Madrer and S. Vaknine-Treidel.

Data used in the preparation of this article were obtained from the Parkinson’s Progression Markers Initiative (PPMI) database (www.ppmi-info.org/access-data-specimens/download-data). For up-to-date information on the study, visit ppmi-info.org. PPMI – a public-private partnership – is funded by the Michael J. Fox Foundation for Parkinson’s Research and funding partners, including 4D Pharma, AbbVie Inc., AcureX Therapeutics, Allergan, Amathus Therapeutics, Aligning Science Across Parkinson’s (ASAP), Avid Radiopharmaceuticals, Bial Biotech, Biogen, BioLegend, Bristol Myers Squibb, Calico Life Sciences LLC, Celgene Corpora-tion, DaCapo Brainscience, Denali Therapeutics, The Edmond J. Safra Foundation, Eli Lilly and Company, GE Healthcare, GlaxoSmithKline, Golub Capital, Handl Therapeutics, Insitro, Janssen Pharmaceuticals, Lundbeck, Merck & Co. Inc., Meso Scale Diagnostics, LLC, Neurocrine Biosci-ences, Pfizer Inc., Piramal Imaging, Prevail Therapeutics, F. Hoffmann-La Roche Ltd and its af-filiated company Genentech Inc., Sanofi Genzyme, Servier, Takeda Pharmaceutical Company, Teva Neuroscience Inc., UCB, Vanqua Bio, Verily Life Sciences, Voyager Therapeutics Inc., and Yu-manity Therapeutics Inc.

